# Corticolimbic connectivity mediates the relationship between pubertal timing and mental health problems

**DOI:** 10.1101/2023.02.13.528385

**Authors:** Nandita Vijayakumar, Sarah Whittle, Timothy J. Silk

**Affiliations:** School of Psychology, Deakin University, Burwood, Victoria, Australia; Melbourne Neuropsychiatry Centre, Department of Psychiatry, The University of Melbourne and Melbourne Health, Victoria, Australia; Developmental Imaging, Murdoch Children’s Research Institute, Parkville, Victoria, Australia

**Keywords:** pubertal timing, connectivity, resting-state, mental health, family environment

## Abstract

**Background:** Undergoing puberty ahead of peers (“earlier pubertal timing”) is an important risk factor for mental health problems during early adolescence. The current study examined pathways between pubertal timing and mental health via connectivity of neural systems implicated in emotional reactivity and regulation (specifically corticolimbic connections) in 9- to 14-year-olds.

**Method:** Research questions were examined in the Adolescent Brain Cognitive Development (ABCD) Study, a large population representative sample in the United States. Linear mixed models examined associations between pubertal timing and resting-state corticolimbic connectivity. Significant connections were examined as potential mediators of the relationship between pubertal timing and mental health (withdrawn depressed and rule-breaking delinquency) problems. Exploratory analyses interrogated whether the family environment moderated neural risk patterns in those undergoing puberty earlier than their peers.

**Results:** Earlier pubertal timing was related to decreased connectivity between limbic structures (bilateral amygdala and right hippocampus) and the cingulo-opercular network (CON), as well as between the left hippocampus and ventral attention network (VAN). Corticolimbic connections also mediated the relationship between earlier pubertal timing and increased withdrawn depressed problems (but not rule-breaking delinquency). Finally, parental acceptance buffered against limbic-CON connectivity patterns that were implicated in withdrawn depressed problems in those undergoing puberty earlier than their peers.

**Conclusion:** Findings highlight the role of decreased corticolimbic connectivity in mediating pathways between earlier pubertal timing and withdrawn depressed problems, and we present preliminary evidence that the family environment may buffer against these neural risk patterns during early adolescence.

---

Undergoing pubertal maturation earlier than one’s peers (referred to as earlier pubertal timing) is consistently identified as a predictor for adolescent mental health problems (Ullsperger & Nikolas, 2017). Adolescence is also characterized by critical development of the brain’s functional networks, with pubertal processes driving certain aspects of maturation (Vijayakumar et al., 2018). In particular, it is often postulated that puberty may play a role in changing connectivity between cortical and limbic structures (Colich & McLaughlin, 2022). Corticolimbic functional connectivity is also commonly implicated in mental health problems (Chahal et al., 2020; Marusak et al., 2016; Schmitt, 2022), and is thought to directly contribute to alterations in emotion regulation. It thus represents a potential pathway between earlier pubertal timing and mental health problems. Understanding this mechanistic pathway will aide the identification of targets for mental health interventions during the adolescent years.

## Puberty and corticolimbic connectivity

During puberty, the release of androgen and gonadal hormones trigger the process of physical and sexual development. These same hormones also act via androgen and estrogen receptors in the brain to modulate neurotransmission and influence synaptic function, myelination, and neurite growth (Piekarski et al., 2017). Hormone receptor density is highest within limbic regions, particularly the amygdala and hippocampus (Kashon & Sisk, 1994; Milner et al., 2010). While receptor density is more sparse in the cortex, recent rodent studies illustrate that pubertal processes also influence cortical development – particularly within the medial PFC (Delevich et al., 2021). Further, the properties of neuronal projections between the amygdala and PFC are also in flux during puberty (Cressman et al., 2010; Cunningham et al., 2002), suggesting that corticolimbic connections are changing.

While a growing literature examines maturation of the human brain during puberty, few studies have focused on connectivity. Those directly interrogating corticolimbic connectivity have noted reductions during resting-state with increasing pubertal stage (conceptualized as the observable physical changes driven by hormones), such as decreased hippocampal-dorsal ACC connectivity in females (van Duijvenvoorde et al., 2019). Similarly, recent investigations in a large representative US sample have reported decreased connectivity between the amygdala and cingulo-opercular network (CON; comprising the dorsal anterior cingulate and insular cortices) at higher pubertal stage (Thijssen et al., 2020). Others have examined task-evoked connectivity, with some identifying similar decreases in amygdala-insula connectivity with increasing androgen hormones (specifically dehydroepiandrosterone [DHEA] in females (Barendse et al., 2020)) and longitudinal decreases in amygdala-OFC connectivity with increasing testosterone levels during threat-processing (Spielberg et al., 2015). However, inconsistences include increased amygdala-dorsal ACC connectivity with testosterone levels during threat-processing (in males, Barendse et al., 2020) and increased amygdala-vmPFC connectivity with faster pubertal tempo (i.e., longitudinal increases in pubertal stage) during emotion processing (Miller et al., 2020). Thus, there is evidence that corticolimbic connectivity – particularly between the amygdala and prefrontal cortex – changes during puberty. However, with the exception of Barendse and colleagues (2020), these studies have not specifically interrogated pubertal timing – the stage of an individual relative to same-age peers. While studies use varying methods of modelling age, it is unclear whether they are capturing individual differences in timing. Therefore, it remains uncertain whether connectivity patterns reflect normative changes through the progression of pubertal stages, or accelerated patterns (beyond the group average) in those with early timing.

## Implications for mental health

Notably, aberrant patterns of corticolimbic connectivity are consistently implicated in mental health problems (Chahal et al., 2020; Marusak et al., 2016; Schmitt, 2022). Many studies have identified altered resting-state amygdala-PFC connectivity in adolescent depression; several highlight decreased connectivity with higher-order cognitive regions (Scheuer et al., 2017; Tang et al., 2018; Wu et al., 2020), but there remain inconsistencies (Brieant et al., 2021). While less research has interrogated externalizing problems, some have implicated similar corticolimbic networks (Kim et al., 2016; Thijssen et al., 2021). Critically, accelerated reductions in amygdala-PFC connectivity is thought to reflect dysregulated top-down control of limbic reactivity to environmental stressors, thus suggesting aberrant emotion processing and regulation (Pannekoek et al., 2014; Perlman et al., 2012). Others propose that accelerated maturation may prematurely terminate sensitive periods of neural plasticity that support the acquisition of emotion regulation skills (Callaghan & Tottenham, 2016). Meta-analyses have also implicated increased connectivity between the amygdala and default-mode network in adolescent depression (Tang et al., 2018), which is thought to reflect recursive self-referential thoughts and negative affective response patterns (Cooney et al., 2010; Cullen et al., 2014). However, some note decreased amygdala connectivity when considering the vmPFC alone – one component of the default mode network (Hanson et al., 2019). Thus, although inconsistencies remain with regards to directionality, findings suggest that altered resting-state corticolimbic connectivity may represent a potential pathway between pubertal timing and mental health problems.

Indeed, prominent models of neurodevelopment propose that pubertal processes drive the maturation of limbic systems early in adolescence, while experiential factors lead to protracted maturation of cortical networks into young adulthood, thus resulting in “normative” patterns of corticolimbic mismatch (i.e., relatively earlier limbic compared to cortical maturation) (Shulman et al., 2016). Accelerated maturation in individuals undergoing puberty earlier than their peers may therefore exacerbate this mismatch, with resultant corticolimbic dysregulation increasing risk for mental health problems. However, few have investigated pathways between pubertal timing, corticolimbic connectivity and mental health. Kircanski and colleagues (2019) showed that accelerated frontolimbic white matter development during early puberty predicted greater depression symptomatology. Spielberg et al. (2015) showed that increased testosterone was related to decreased amygdala-OFC functional connectivity during threat processing, which was in turn related to increased withdrawn temperament. Barendse et al. (2020) found similar patterns predicting anxiety in adolescent females, but opposing effects in males. To our knowledge, the only study to examine resting-state connectivity failed to identify any associations between puberty-related network changes and internalizing problems, although it did not focus on corticolimbic connections (Ernst et al., 2019).

## Contextual amplification

models of puberty propose that early timing may be considered a biological marker of vulnerability that interacts with environmental stressors to predict mental health problems. For example, some research suggests that early timing is only related to depression and conduct problems in the context of adverse family environments (Rudolph & Troop-Gordon, 2010). Further, Vijayakumar and colleagues (2022) showed that supportive environments act to buffer against associations between timing and depressive symptoms and delinquent behavior problems. While unfavourable social environments may exacerbate the stress of undergoing puberty earlier than peers, with multiple concurrent challenges overtaxing under-developed cognitive and emotional resources and leading to mental health problems, favourable social contexts may better support individuals to navigate through the challenges of puberty without adverse consequences for their mental health. Thus, social contexts may play a role in moderating corticolimbic connectivity that is implicated in mental health problems among those experiencing puberty earlier than peers. A growing literature highlights the potential for social contexts – particularly family environments (Pozzi et al., 2021; Thijssen et al., 2017, 2020) – to influence connectivity, and there is preliminary evidence that parenting may moderate associations between pubertal timing and vlPFC function during threat processing (Barbosa et al., 2018). Identifying such social contexts can inform intervention targets for those at heightened vulnerability during the early adolescent years.

## The current study

The goal of the current study was to investigate associations between pubertal timing, corticolimbic connectivity and mental health problems. We focused on the limbic structures of the amygdala and hippocampus given the density of pubertal hormone receptors in these regions. We examined their connectivity to large scale, distributed functional networks in the cortex during resting-state, building upon research that has implicated these networks in mental health problems. Associations were investigated in the ABCD Study, a large population representative sample in the United States. The first aim was to examine associations between pubertal timing and corticolimbic connectivity, hypothesizing that earlier timing would be related to decreased connectivity between amygdala/hippocampus and higher-order cognitive networks (e.g., cingulo-opercular, frontoparietal and default mode). Next, we examined whether functional connectivity mediated the well-characterized association between pubertal timing and mental health problems, focusing on depression and rule-breaking delinquency that have been most consistently implicated in the literature (Vijayakumar & Whittle, 2022). It was hypothesized that decreased connectivity of the amygdala/hippocampus and higher-order networks would significantly mediate the association between earlier timing and greater mental health problems. Finally, an exploratory aim was to identify family environments that moderate the functional connectivity patterns implicated in mental health problems. We hypothesized that positive family environments (characterized by either high levels of parental acceptance or low levels of family conflict) would buffer patterns of functional connectivity that were related to mental health problems (e.g., positive family environments would lead to less reductions in connectivity in adolescents with early timing).

## Methods

The sample was derived from the ABCD Study (Release 4.0 downloaded from the National Institute of Mental Health Data Archive’s ABCD Collection). Analyses focused on data from baseline and 2-year follow-up assessments when participants were 9-11 and 10.5-13.5 years old, respectively. Data points were excluded in the following successive order: missing puberty (N = 600), mental health (N = 2243) or rsfMRI data (N = 1659), as well as rsfMRI data recommended for exclusion (described further in below, N = 2637). This resulted in a final sample of 15197 data points (7400 females) from 10228 participants (5000 females), including 8689 unique families (and 1496 sibling sets). Data was collected across 22 sites in the United States. The sample comprised 52.9% White, 19.8% Hispanic, 14.7% Black/African American, and 12.2% Other/Mixed races. Refer to Table 1 for further descriptive information.

**Table 1.** Sample characteristics

|  | Females |  |  |  |  |  |  |  | Males |  |  |  |  |  |  |  |
| --- | --- | --- | --- | --- | --- | --- | --- | --- | --- | --- | --- | --- | --- | --- | --- | --- |
|  | Baseline |  |  |  | 2-year follow-up |  |  |  | Baseline |  |  |  | 2-year follow-up |  |  |  |
|  | Mean | SD | Min | Max | Mean | SD | Min | Max | Mean | SD | Min | Max | Mean | SD | Min | Max |
| Age | 9.92 | 0.62 | 8.92 | 11.00 | 11.89 | 0.64 | 10.67 | 13.83 | 9.96 | 0.63 | 8.92 | 11.08 | 11.95 | 0.64 | 10.58 | 13.58 |
| PDS | 1.70 | 0.54 | 1.00 | 4.00 | 2.34 | 0.66 | 1.00 | 4.00 | 1.64 | 0.49 | 1.00 | 4.00 | 1.85 | 0.54 | 1.00 | 4.00 |
| CBCL depression | 44.73 | 9.81 | 34.00 | 82.00 | 43.74 | 9.48 | 34.00 | 82.00 | 46.04 | 10.56 | 33.00 | 84.00 | 44.89 | 9.94 | 33.00 | 80.00 |
| CBCL delinquency | 47.46 | 10.53 | 33.00 | 93.00 | 47.23 | 10.43 | 33.00 | 86.00 | 49.04 | 10.55 | 34.00 | 88.00 | 47.93 | 10.54 | 34.00 | 90.00 |
| CRPBI | 2.80 | 0.29 | 1.00 | 3.00 | 2.83 | 0.27 | 1.20 | 3.00 | 2.77 | 0.30 | 1.00 | 3.00 | 2.80 | 0.29 | 1.00 | 3.00 |
| FES | 1.89 | 1.91 | 0.00 | 9.00 | 1.82 | 1.83 | 0.00 | 9.00 | 2.10 | 1.95 | 0.00 | 9.00 | 1.89 | 1.79 | 0.00 | 9.00 |
| INR | 4.02 | 2.87 | 0.03 | 15.39 | 4.09 | 2.77 | 0.05 | 14.78 | 4.04 | 2.83 | 0.03 | 15.39 | 4.22 | 2.81 | 0.06 | 14.78 |

CBCL = Child Behavior Checklist, CRPBI = Children's Report of Parental Behavior Inventory, FES = Family Environment Scale, INR = Income-to-need ratio
(calculation outlined in Supplementary Material), PDS = Pubertal Development Scale.

Ethical review and approval of ABCD’s protocol, as well as informed consent procedures, are outlined by Clark et al. (2017). The use of pre-existing and non-identifiable data in the current analyses was approved by Deakin University’s Human Research Ethics Committee.

### Measures

#### Pubertal timing

Adolescents completed the Pubertal Development Scale (PDS, Petersen et al., 1988) consisting of five questions that assess height growth, body hair and skin changes, as well as breast development and menarche in females, and facial hair and voice changes in males. Items were rated on a 4-point scale and mean scores were calculated across items. Pubertal timing was calculated within each sex by regressing age from pubertal stage using linear mixed model: PDS ∼ age + age^2^ + (1|subject_id) + (1|family_id) + (1| site_id). A quadratic (fixed) effect of age was modelled given nonlinear trends identified in prior literature (Vijayakumar et al., 2021; Wierenga et al., 2018), and random effects of subject, family and site IDs accounted for the nested nature of the data. Model residuals were saved as an index of pubertal stage relative to same-age and -sex peers. Positive and negative values reflect earlier and later pubertal timing, respectively. Refer to Figure S1 for an illustration of the association between pubertal stage and age, as well as the relative distribution of pubertal timing (i.e., model residuals).

#### rsfMRI

Adolescents underwent MRI scans in a 3T scanner (Siemens, Philips, or General Electric). They completed four or five resting-state MRI scans (lasting 5 minutes), with their eyes open and fixated on a crosshair (further details provided by Casey et al., 2018). Pre-processing was undertaken by the ABCD Data Analysis and Informatics Core using a standardized pipeline. Pre-processing involved removal of initial frames, normalization, regression and temporal filtering (Hagler et al., 2019). Time courses were then projected onto each participant’s cortical surface, and average cortical time courses were calculated for 12 resting state networks defined by the Gordon parcellation scheme (auditory network [AN], cingulo-opercular network [CON], cinguloparietal network [CPN], dorsal attention network [DAN], default mode network [DMN], frontoparietal network [FPN], retrosplenial temporal network [RTN], sensorimotor [hand] network [SMN-H], sensorimotor [mother] network [SMN-M], salience network [SN], ventral attention network [VAN] and visual network [VN]). Average time courses were also calculated for FreeSurfer-defined amygdala and hippocampus (separately for each hemisphere). Correlations between limbic ROIs and cortical networks were calculated and Fisher-Z transformed, providing summary measures of cortico-limbic connectivity. The current study utilized these indices of connectivity via the ABCD Study’s tabulated data. As recommended (Hagler et al., 2019), data from the tabulated output was excluded if it failed to pass FreeSurfer quality control (fsqc_qc was 0), had excessive motion (rsfmri_c_ngd_ntpoints less than 375), or incidental clinical MRI findings (mrif_score greater than 2). Two additional motion metrics were included as covariates in analyses: mean framewise displacement (motion) and outlier count (rsfmri_c_ngd_nvols - rsfmri_c_ngd_ntpoints).

#### Mental health problems

Parents completed the Child Behavior Checklist for Ages 6-18 (CBCL/6-18). Analyses focused on withdrawn depression and rule-breaking delinquency syndrome T-scores. To deal with positive skew, variables were re-centered by subtracting the minimum value, and Poisson distribution was used in statistical modelling.

#### Family environment

Adolescents completed the Children’s Report of Parental Behavior Inventory (CRPBI) and Family Environment Scale (FES). CRPBI includes a parental acceptance subscale comprising 5 items (rated on a 3-point Likert Scale), which was completed about the primary caregiver. FES includes a subscale on family conflict comprising 9 items about members of the family unit (e.g., “family members often criticize each other”). For both scales, items are summed, with higher scores reflecting greater parental acceptance and family conflict, respectively.

### Statistical analyses

#### Puberty and corticolimbic connectivity

Linear mixed models examined whether corticolimbic connectivity differed across pubertal timing (i.e., connectivity ∼ timing), using *lme4* (Bates et al., 2015) in R. While pubertal stage was not a focus on this study, we also ran models that examined associations between connectivity and pubertal stage (i.e., PDS score) to aide interpretation of findings related to pubertal timing. Separate models were conducted for each connectivity variable (N = 48). Covariates included the fixed effects of sex, mean framewise displacement, outlier count, and participation following the start of the COVID19 pandemic (yes/no). Random effects of subject, family and site IDs accounted for the nested nature of the data. To ensure that findings were robust, we undertook a within sample split-half replication; our sample was split into discovery and replication datasets (based on family and subject clustering to ensure independence, using “groupKfold” in *caret* (Kuhn, 2008)), and mixed models were conducted on each dataset. We controlled for multiple comparisons at a false discovery rate (FDR) of 5% in each of the discovery and replication dataset, and only findings surviving FDR correction in both datasets were considered significant.

Supplemental analyses examined associations between corticolimbic connectivity and pubertal timing when incorporating additional covariates of income-to-needs ratio (INR; calculation outlined in the Supplement) and race/ethnicity. These variables were not included in primary models to avoid further loss of sample size due to missingness (10% of the sample [n=1450] were missing one or more variables). We also explored whether associations between corticolimbic connectivity and pubertal timing differed across sex (connectivity ∼ timing*sex) and age (connectivity ∼ timing*age) but failed to identify any significant effects (see Table S3).

#### Pubertal timing, corticolimbic connectivity, and mental health

For significant connections, we examined potential mediation of the association between timing and mental health problems via corticolimbic connectivity. We undertook mediation analyses through Bayesian modelling with *brms* (Bürkner, 2017) which supports multilevel models with multiple random effects. Modelling involved calculation of 95% credible intervals (CIs) to gauge uncertainty of indirect estimates, using Markov chain Monte Carlo simulations. Given the observed data, there is 95% probability of identifying the true estimate within the credible interval. We calculated CIs based on highest density intervals, which always include the mode of posterior distributions. As we did not have prior knowledge of anticipated mediation effects, we used weakly informative priors that decrease the likelihood of estimating unrealistically small or large effects, without substantively influencing regression parameters (Gelman et al., 2008). We used the default number of 4 Markov chains but increased sampling iterations to 10000 (with 2000 warmup) as chains did not adequately converge when using the default value of 2000.

Separate mediation models examined each combination of connectivity and mental health problems (withdrawn depression and rule-breaking delinquency). For each model, a multivariate framework specified both paths A and B, with the same random effects and covariates as the linear mixed effects models. Path A was modelled with the (default) Gaussian distribution, while Path B was modelled with a Poisson distribution due to non-normality of the CBCL variables. Model convergence was examined based on Rhat values (< 1.05) and posterior predictive checks. Next, the indirect effect was calculated using the product of coefficients method, by multiplying the regression coefficient for path A and B in the posterior distribution to derive a distribution for the mediation effect (similar to bootstrapped mediation). Significant mediation was determined when the 95% CI range (reflecting the highest density interval across posterior distributions) did not include zero.

#### Moderation by the family environment

Exploratory models examined whether the family environment moderated associations between pubertal timing and corticolimbic connectivity. For connectivity variables identified as significant mediators, linear mixed models (with *lme4*) were used to examine whether parental acceptance or family conflict interacted with pubertal timing to predict corticolimbic connectivity (connectivity ∼ pubertal timing*family environment), with the same covariates and random effects as prior models. Finally, moderated mediation models examined whether the family environment moderated indirect pathways between timing and mental health problems. Specifically, models for path A and B incorporated a main effect of family environment and an interaction term with pubertal timing (separately for parental acceptance and family conflict). Moderation effects were further interrogated by calculation of mediation pathways at high (+1SD above mean) and low (−1SD below mean) levels of the family environment.

## Results

### Puberty and corticolimbic connectivity

Patterns of resting-state corticolimbic connectivity across pubertal stage and timing are illustrated in Figure 1 (with statistics presented in Table S2). More advanced pubertal stage was predominantly related to decreased connectivity across corticolimbic networks, with strongest effects present for left amygdala-CON and left hippocampus-VAN. Comparatively, increased connectivity was evident for limbic-AN and a few other connections. Earlier pubertal timing was associated with decreased functional connectivity between limbic structures (bilateral amygdala and right hippocampus) and CON, as well as between the left hippocampus and VAN. These associations remained significant when controlling for race/ethnicity and INR, aside from connectivity of the right hippocampus and CON (refer to Supplement).

**Figure 1.**
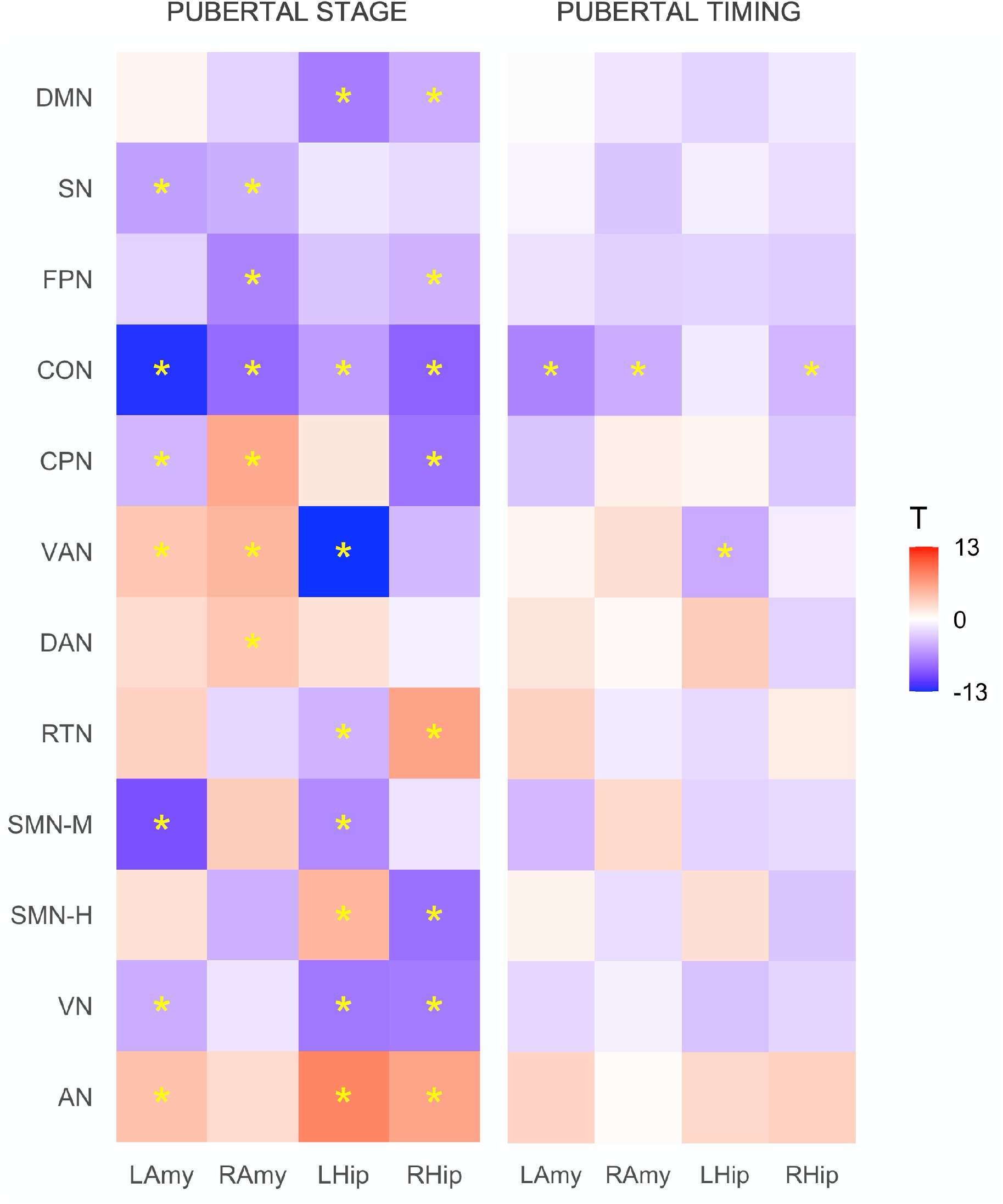
Associations between resting-state corticolimbic connectivity and i) pubertal stage and ii) pubertal timing. Heatmaps represent T-statistics (T) of models on the full dataset, while significance (*) is based on FDR 0.05 for both discovery and replication sets. LAmy = Left Amygdala, RAmy = Right Amygdala, LHip = Left Hippocampus, RHip = Right Hippocampus, DMN = Default Mode Network, SN = Salience Network; FPN = Frontoparietal Network, CON = Cingulo-Opercular Network, CPN = Cingulo-Parietal Network, VAN = Ventral Attention Network, DAN = Dorsal Attention Network, RSN = Retrosplenial Temporal Network, SMN-M = Somatomotor Network – Mouth, SMN-H = Somatomotor Network – Hand, VN = Visual Network, AN = Auditory Network

### Pubertal timing, corticolimbic connectivity, and mental health

Mediation analyses examined whether the four corticolimbic connections that were significantly related to pubertal timing mediated pathways to mental health problems (statistics from all multivariate models are presented in Table S4). As illustrated in Figure 2, analyses revealed a direct effect of pubertal timing on withdrawn depression, as well as an indirect effect mediated via amygdala-CON connectivity (accounting for up to 7% of the total effect). Specifically, decreased connectivity (of left amygdala-CON, right amygdala-CON, right hippocampus-CON and left hippocampus-VAN) mediated the relationship between earlier pubertal timing and higher levels of withdrawn depression. There was also a direct effect of pubertal timing on rule-breaking delinquency, with earlier timing predicting higher levels of delinquency, and there was an indirect pathway between timing and delinquency via right hippocampus-CON connectivity. There were no further indirect pathways involving any of the other corticolimbic connections. Most of these associations remained significant when controlling for race/ethnicity and INR, aside from those involving the right hippocampus and CON (refer to Supplement).

**Figure 2.**
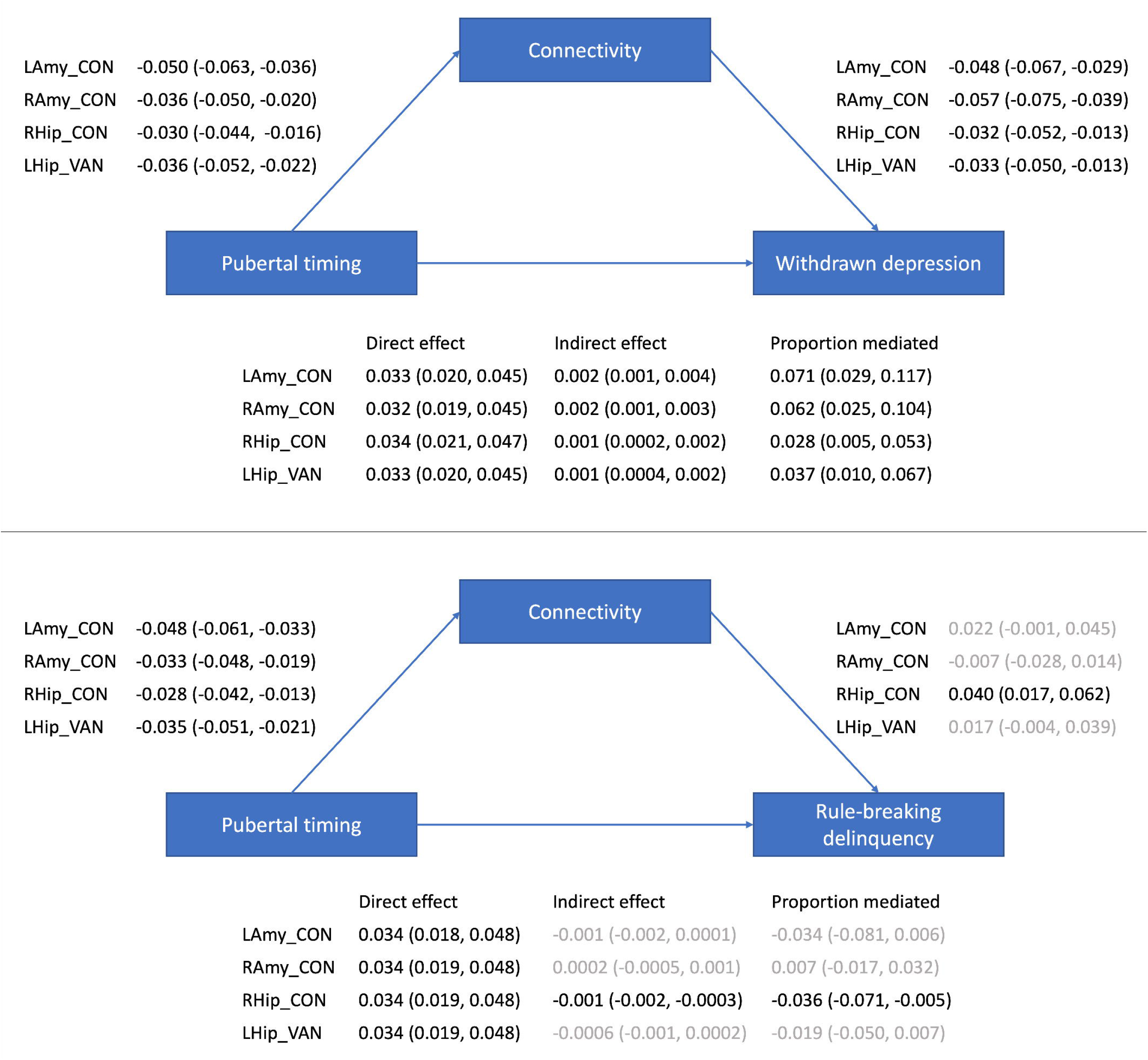
Mediation models examining indirect pathways between pubertal timing and mental health problems via resting-state corticolimbic connectivity (estimates and 95% CIs presented). LAmy = Left Amygdala, RAmy = Right Amygdala, LHip = Left Hippocampus, RHip = Right Hippocampus, CON = Cingulo-Opercular Network, VAN = Ventral Attention Network

### Pubertal timing, family environment, and corticolimbic connectivity

Exploratory analyses failed to identify significant moderation between pubertal timing and family conflict in predicting connectivity, but significant moderation was identified for parental acceptance across left amygdala-CON, right amygdala-CON and right hippocampus-CON (but not left hippocampus-VAN; see Table 2). As illustrated in Figure 3, high levels of parental acceptance buffered against reductions in connectivity, while low levels increased reductions in connectivity, in those with earlier timing. Further, there was significant moderation of indirect pathways between pubertal timing and withdrawn depression by parental acceptance via left amygdala-CON (−0.004 [CIs: -0.007, -0.001]), right amygdala-CON (−0.004 [CIs: -0.007, -0.001]) and right hippocampus-CON (−0.002 [CIs: -0.004, - 0.0001]), as well as between pubertal timing and rule-breaking delinquency by right hippocampus-CON (0.003 [CIs: 0.0003, 0.005]). For all models, the indirect pathway was stronger at low levels of parental acceptance and not present at high levels of parental acceptance.

**Table 2.** Moderation of pubertal timing and the family environment predicts corticolimbic connectivity

|  | Family conflict | Parental acceptance |
| --- | --- | --- |
| LAmy-CON | B = 0.000, SE = 0.001,<br>T = -0.305, p = 0.761 | B = 0.021, SE = 0.007,<br>T = 3.082, p = 0.002 |
| RAmy-CON | B = -0.000, SE = 0.001,<br>T = -0.672, p = 0.501 | B = 0.021, SE = 0.007,<br>T = 2.877, p = 0.005 |
| RHip-CON | B = -0.000, SE = 0.001,<br>T = -0.550, p = 0.582 | B = 0.018, SE = 0.006,<br>T = 2.832, p = 0.005 |
| LHip-VAN | B = -0.000, SE = 0.001,<br>T = -0.266, p = 0.790 | B = 0.002, SE = 0.007,<br>T = 0.219, p = 0.827 |

Statistics are presented for the interaction term between pubertal timing and family environment predicting connectivity. LAmy = Left Amygdala, RAmy = Right Amygdala, LHip = Left Hippocampus, RHip = Right Hippocampus, CON = Cingulo-Opercular Network, VAN = Ventral Attention Network

**Figure 3.**
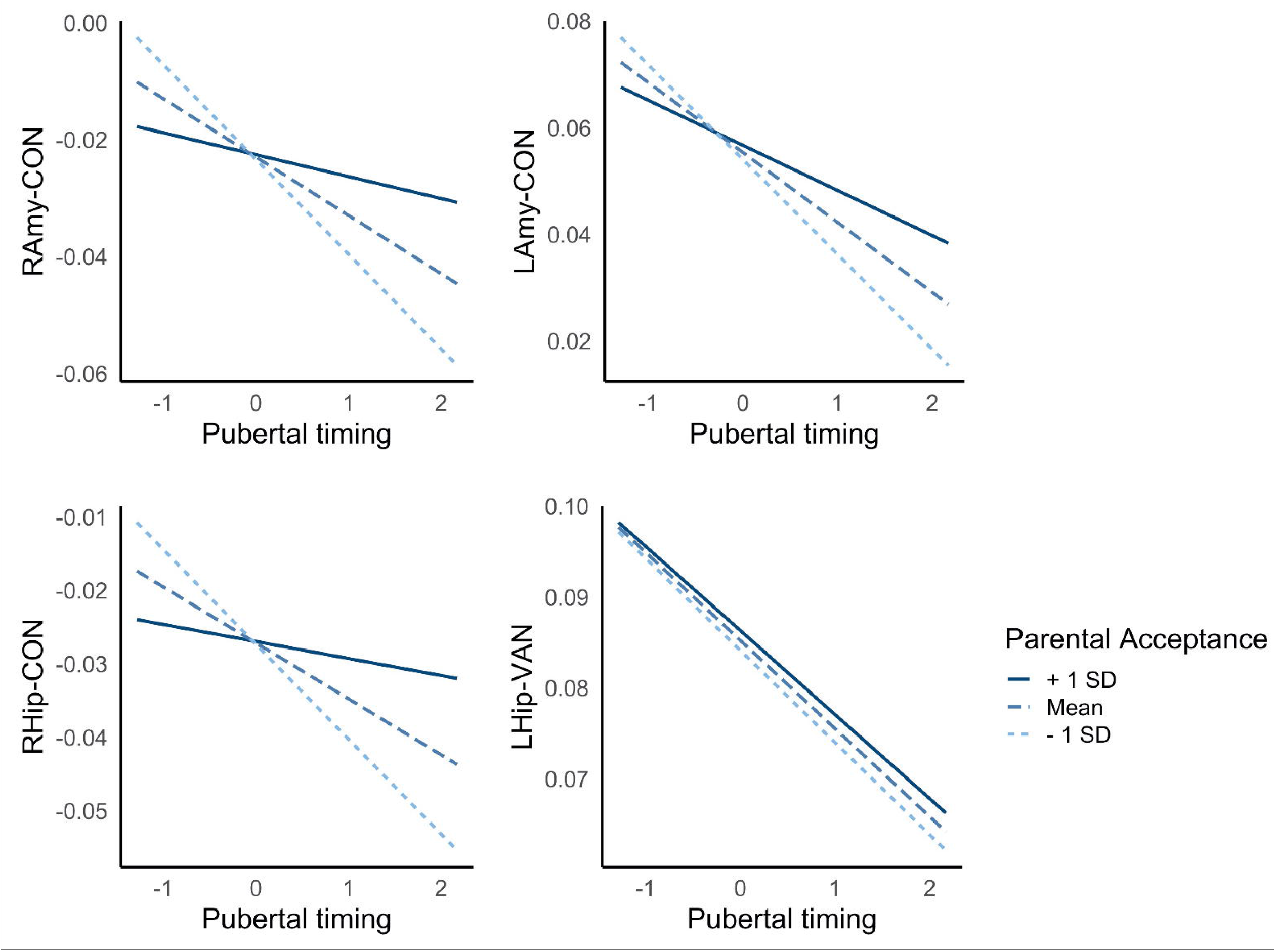
Interactions between pubertal timing and parental acceptance predict resting-state amygdala-CON connectivity. LAmy = Left Amygdala, RAmy = Right Amygdala, CON = Cingulo-Opercular Network

## Discussion

The current study investigated the role of corticolimbic connectivity in mediating well-established associations between earlier pubertal timing and mental health problems. Findings highlighted decreased corticolimbic connectivity in adolescents undergoing puberty earlier than their peers, which mediated associations between earlier timing and withdrawn depressed problems. Further, a positive family environment characterized by parental acceptance was found to buffer against these risk patterns, as it was related to less reductions in connectivity in those with earlier pubertal timing.

There was a general decrease in corticolimbic connectivity in adolescents at later stages of puberty, particularly for higher order networks (i.e., CON, frontoparietal, default mode and salience networks). These findings extend prior literature (Thijssen et al., 2020; van Duijvenvoorde et al., 2019) to highlight widespread resting-state connectivity changes between limbic regions (specifically amygdala and hippocampus) and most cortical networks during puberty, reaching well beyond the prefrontal cortex. Findings are broadly similar to age-related changes that have been identified during this period (van Duijvenvoorde et al., 2019) which is expected given the strong correlation between age and pubertal stage. However, as there are inconsistencies in previously identified patterns of corticolimbic changes in prior literature, the current findings help to aide interpretation of the primary investigations into pubertal timing.

Beyond group-level changes during puberty, those who underwent puberty earlier than their peers exhibited greater (than average) reductions in bilateral amygdala-CON, right hippocampus-CON and left hippocampus-VAN connectivity. These findings provide strong support for involvement of the CON, which is implicated in cognitive control processes such as maintenance of alertness and task-set over time (Menon & D’Esposito, 2022). Animal studies have reported high density of hormone receptors in the amygdala and hippocampus, as well as extensive anatomical connections between the amygdala and frontal and insular cortices (Aggleton et al., 1980; Ghashghaei et al., 2007). Overall, the pattern of effects is suggestive of accelerated biological development in those who undergo pubertal earlier than peers, when compared to connectivity changes identified for normative progression through pubertal stage. Indeed, both earlier pubertal timing and decreased corticolimbic connectivity have been independently considered markers of accelerated biological development, particularly within contexts of early life stress (Colich & McLaughlin, 2022). Earlier timing supports reproductive success in conditions of adversity and uncertainty (Belsky, 2019). Decreased connectivity (particularly amygdala-PFC) may also reflect accelerated maturation of emotional systems in response to external stressors (increasing top-down signalling through activation of higher-order cognitive control networks (Gee et al., 2013)). Thus it is possible that stress-induced changes in the hypothalamic-pituitary-adrenal system and downstream changes in the hypothalamic-pituitary-gonadal system may underlie associations between early timing and greater-than-normative reductions in corticolimbic connectivity (Colich & McLaughlin, 2022).

There is a well-established relationship between early pubertal timing and mental health problems, particularly depression and conduct/delinquency problems (Ullsperger & Nikolas, 2017). It is often theorized that altered corticolimbic connectivity may mediate this association (Colich & McLaughlin, 2022), but few have empirically investigated this relationship. The current findings provide support for this model, based on significant indirect pathways between early timing and mental health problems via decreased corticolimbic connectivity. In particular, effects for withdrawn depression were robust to the potential confounding effects of demographic characteristics (race/ethnicity and socioeconomic status). Findings broadly align with prior literature highlighting a potential role of accelerated corticolimbic connectivity during threat processing (Barendse et al., 2020; Spielberg et al., 2015) and similar patterns of white matter development (Kircanski et al., 2019) in predicting internalizing problems for those with advanced pubertal maturation. While the only prior study of resting-state functional connectivity failed to identify such pathways, it did not specifically focus on corticolimbic connections (Ernst et al., 2019). The prevalence of hormone receptors in the amygdala, and its white matter connections with the frontal and insular cortex, may place amygdala-CON connectivity as key risk pathways in those with earlier puberty. While decreased corticolimbic connectivity may reflect earlier maturation of emotional systems in response to early life stress, it may also decrease the window of neural plasticity to support maturation of emotional regulatory circuits and thus increase risk for depression (Callaghan & Tottenham, 2016).

Finally, there were significant interactions between the family environment and pubertal timing in predicting amygdala-CON connectivity; low levels of parental acceptance exacerbated the negative relationship between timing and connectivity, while high levels buffered against this pattern. Further, there was some support for contextual amplification models as parental acceptance moderated indirect pathways between pubertal timing and withdrawn depression via amygdala-CON connectivity, such that indirect pathways were not present at high levels of parental acceptance. This finding adds to the only prior study to examine and identify interactions between pubertal timing and parenting in relation to adolescent neural functioning (Barbosa et al., 2018). Individuals undergoing earlier puberty lean on their parents for guidance as they face the social challenges of this period (such as being physically different to their peers), and positive family environments are likely to be better equipped to support youth to navigate through these stressors (Rudolph & Troop-Gordon, 2010). Thus, to the extent that negative amygdala-CON connectivity reflects early activation of emotional circuits to respond to external stressors (Callaghan & Tottenham, 2016), we speculate that those in positive family environments may also receive external support with emotion regulation and may therefore require less reductions in connectivity (i.e., less acceleration of biological development) in response to socioemotional demands.

The current study represents one of the few empirical examinations of pathways between earlier pubertal timing, corticolimbic connectivity and mental health problems. It is also the first to specifically examine these relationships based on resting-state connectivity. However, an important limitation is that pubertal timing and stage are highly confounded, and thus is it difficult to tease apart neural changes that are specifically related to early timing. Nonetheless, we characterise patterns of normative neural changes as adolescents progress through the stages of puberty, and additional changes in those with earlier timing. Second, the availability of only two waves of assessments limits modelling of within-subject changes, and thus findings are primarily reflective of between subject effects. Future studies should examine additional waves of the ABCD to investigate longitudinal trajectories, which will also enable disentangling of pubertal timing and tempo. Third, there are many potential roles of the family environment in relation to puberty and corticolimbic connectivity. In addition to its modulation of pathways, there is support for negative family contexts being a sequelae and consequence of early pubertal timing. Indeed, prior research on the ABCD has identified indirect pathways between negative family environments and corticolimbic connectivity via earlier timing (Thijssen et al., 2020). Future studies may combine and compare these models to better understand which components of the family environment act along the pathway to altered corticolimbic connectivity and mental health problems.

In conclusion, the current investigation of the ABCD cohort highlights neural mechanisms that partially account for the increased risk for depression in adolescents experiencing puberty earlier than their peers. In particular, findings implicated decreased connectivity between the amygdala and the cingulo-opercular network involved in higher-order cognitive control. Exploratory analyses provided preliminary support for the role of positive family environments in buffering against these neural risk patterns in adolescents undergoing earlier pubertal maturation, highlighting protective pathways that represent potential intervention targets for those at heightened vulnerability during the early adolescent years.

## Acknowledgements

Data used in the preparation of this article were obtained from the ABCD Study held in the National Institute of Mental Health Data Archive. A full list of supporters is available at https://abcdstudy.org/nihcollaborators. A listing of participating sites and a complete listing of the study investigators can be found at https://abcdstudy.org/principal-investigators.html. ABCD consortium investigators designed and implemented the study and/or provided data but did not participate in analysis or writing of this report.

## Notes

**Conflict of interest:** The authors do not have any conflicts of interest to disclose.

### Competing Interest Statement

The authors have declared no competing interest.

