## Supplementary Material for "Corticolimbic connectivity mediates the relationship between pubertal timing and mental health problems"

**Abbreviations**

LAmy = Left Amygdala, RAmy = Right Amygdala, LHip = Left Hippocampus, RHip = Right Hippocampus, DMN = Default Mode Network, SN = Salience Network; FPN = Frontoparietal Network, CON = Cingulo-Opercular Network, CPN = Cingulo-Parietal Network, VAN = Ventral Attention Network, DAN = Dorsal Attention Network, RSN = Retrosplenial Temporal Network, SMN-M = Somatomotor Network – Mouth, SMN-H = Somatomotor Network – Hand, VN = Visual Network, AN = Auditory Network

**Variable details**

*Table S1.* Variable details

| Variables | ABCD Survey | Questionnaire | Variable Name |
| --- | --- | --- | --- |
| PDS | ABCD Youth Pubertal Development Scale and Menstrual Cycle Survey History | Pubertal Development Scale | pds_ht2_y, pds_skin2_y, pds_bdyhair_y, pds_f4_2_y, pds_f5_y, pds_m4_y, pds_m5_y |
| Withdrawn/Depression | ABCD Parent Child Behavior Checklist Scores | Child Behavior Checklist | cbcl_scr_syn_withdep_t |
| Rule-Breaking Delinquency |  | Child Behavior Checklist | cbcl_scr_syn_rulebreak_t |
| Family Conflict | ABCD Sum Scores Culture & Environment Youth | Family Environment Scale | fes_y_ss_fc_pr |
| Parent Acceptance |  | Child's Report of Parental Behavior Inventory | crpbi_y_ss_parent |
| INR* | ABCD Longitudinal Parent Demographics Survey |  | demo_comb_income_v2_l |

* To calculate the income-to-needs ratio (INR), the categorical household income variable (demo_comb_income_v2_l) was first recoded such that each level referred to the mean of its income bracket. Next, this mean income level was divided by the US Federal Poverty Line based on the number of individuals residing in the house (demo_roster_v2_l). The poverty line for 2017 and 2019 were used for baseline and 2-year follow-up calculations of INR (when the majority of the sample completed assessments at each respective wave).

**Calculation of pubertal timing**

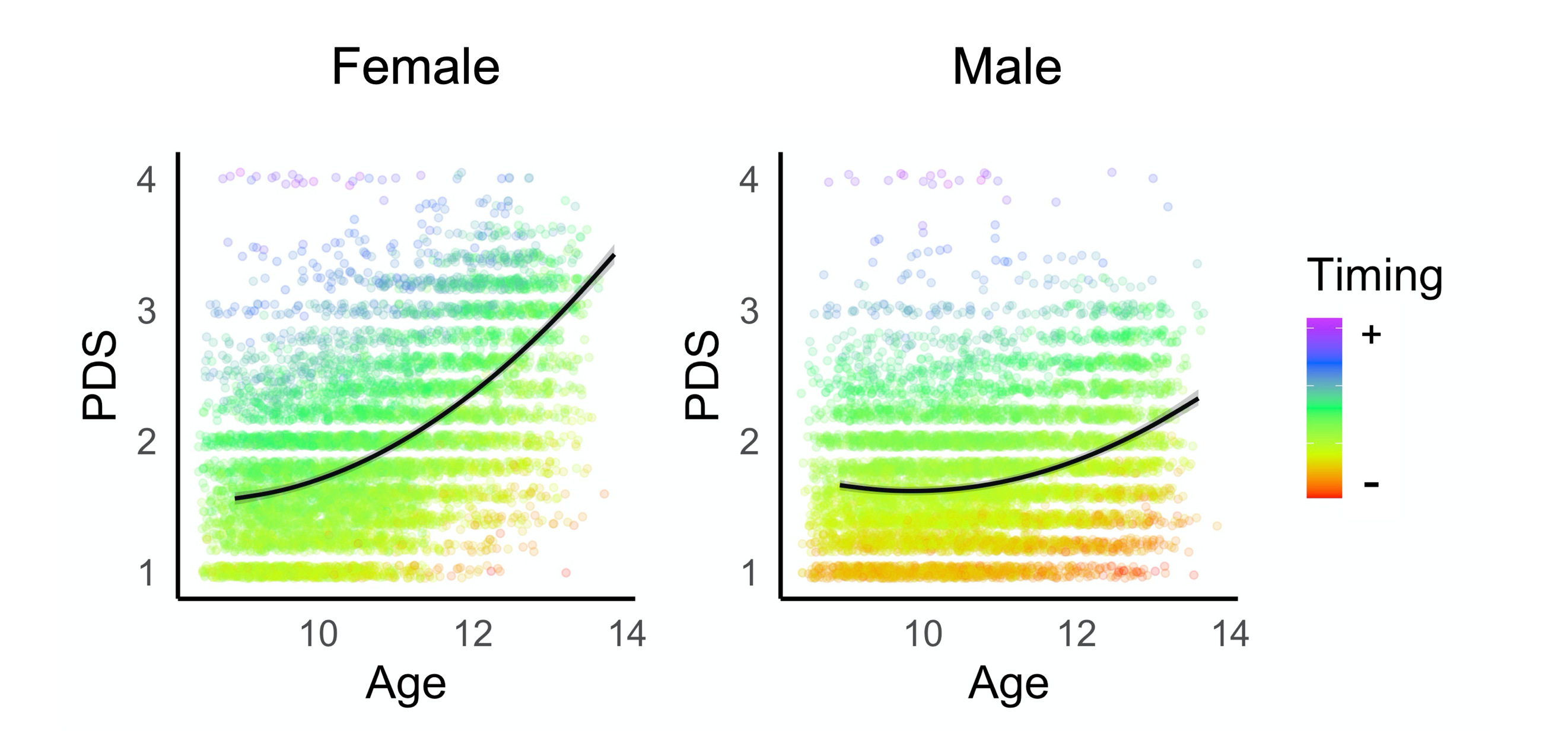

*Figure S1*. Relationship between age and pubertal stage (PDS) in females and males. Colours represent the distribution of pubertal timing (A; positive values reflect earlier timing, while negative values reflect later timing).

**Results tables**

Table S2. *Associations between puberty and corticolimbic connectivity*

| **ROI** | **Network** | **Pubertal Stage** | **Pubertal Timing** |
| --- | --- | --- | --- |
| L_Amy | AN | **B=0.006, SE=0.001, T=4.231, p=0.000** | B=0.006, SE=0.002, T=2.906, p=0.004 |
| L_Amy | VN | **B=-0.005, SE=0.001, T=-4.629, p=0.000** | B=-0.003, SE=0.002, T=-2.191, p=0.028 |
| L_Amy | SMN-H | B=0.002, SE=0.001, T=2.097, p=0.036 | B=0.001, SE=0.001, T=0.955, p=0.340 |
| L_Amy | SMN-M | **B=-0.015, SE=0.002, T=-9.833, p=0.000** | B=-0.009, SE=0.002, T=-3.930, p=0.000 |
| L_Amy | RTN | B=0.004, SE=0.001, T=3.068, p=0.002 | B=0.007, SE=0.002, T=3.146, p=0.002 |
| L_Amy | DAN | B=0.002, SE=0.001, T=2.527, p=0.012 | B=0.002, SE=0.001, T=1.761, p=0.078 |
| L_Amy | VAN | **B=0.004, SE=0.001, T=3.860, p=0.000** | B=0.001, SE=0.002, T=0.690, p=0.490 |
| L_Amy | CPN | **B=-0.002, SE=0.001, T=-4.132, p=0.000** | B=-0.003, SE=0.001, T=-3.287, p=0.001 |
| L_Amy | CON | **B=-0.017, SE=0.001, T=-12.670, p=0.000** | **B=-0.014, SE=0.002, T=-6.910, p=0.000** |
| L_Amy | FPN | B=-0.002, SE=0.001, T=-2.540, p=0.011 | B=-0.002, SE=0.001, T=-1.603, p=0.109 |
| L_Amy | SN | **B=-0.009, SE=0.002, T=-5.379, p=0.000** | B=-0.002, SE=0.002, T=-0.634, p=0.526 |
| L_Amy | DMN | B=0.001, SE=0.001, T=0.713, p=0.476 | B=0.000, SE=0.002, T=0.179, p=0.858 |
| R_Amy | AN | B=0.002, SE=0.001, T=2.397, p=0.017 | B=0.000, SE=0.001, T=0.284, p=0.777 |
| R_Amy | VN | B=-0.002, SE=0.001, T=-1.528, p=0.127 | B=-0.002, SE=0.002, T=-0.827, p=0.408 |
| R_Amy | SMN-H | B=-0.006, SE=0.001, T=-4.524, p=0.000 | B=-0.003, SE=0.002, T=-1.924, p=0.054 |
| R_Amy | SMN-M | B=0.003, SE=0.001, T=3.391, p=0.001 | B=0.003, SE=0.001, T=2.594, p=0.009 |
| R_Amy | RTN | B=-0.001, SE=0.001, T=-2.275, p=0.023 | B=-0.001, SE=0.001, T=-1.181, p=0.238 |
| R_Amy | DAN | **B=0.004, SE=0.001, T=3.859, p=0.000** | B=0.001, SE=0.002, T=0.441, p=0.660 |
| R_Amy | VAN | **B=0.004, SE=0.001, T=4.919, p=0.000** | B=0.003, SE=0.001, T=2.254, p=0.024 |
| R_Amy | CPN | **B=0.008, SE=0.001, T=5.811, p=0.000** | B=0.002, SE=0.002, T=1.073, p=0.283 |
| R_Amy | CON | **B=-0.012, SE=0.001, T=-8.384, p=0.000** | **B=-0.010, SE=0.002, T=-4.606, p=0.000** |
| R_Amy | FPN | **B=-0.009, SE=0.001, T=-6.890, p=0.000** | B=-0.005, SE=0.002, T=-2.615, p=0.009 |
| R_Amy | SN | **B=-0.004, SE=0.001, T=-4.472, p=0.000** | B=-0.005, SE=0.001, T=-3.291, p=0.001 |
| R_Amy | DMN | B=-0.003, SE=0.001, T=-2.564, p=0.010 | B=-0.002, SE=0.002, T=-1.435, p=0.151 |
| L_Hip | AN | **B=0.005, SE=0.001, T=8.072, p=0.000** | B=0.002, SE=0.001, T=2.718, p=0.007 |
| L_Hip | VN | **B=-0.010, SE=0.001, T=-7.735, p=0.000** | B=-0.006, SE=0.002, T=-3.466, p=0.001 |
| L_Hip | SMN-H | **B=0.008, SE=0.002, T=4.971, p=0.000** | B=0.005, SE=0.002, T=2.164, p=0.030 |
| L_Hip | SMN-M | **B=-0.008, SE=0.001, T=-6.545, p=0.000** | B=-0.004, SE=0.002, T=-2.474, p=0.013 |
| L_Hip | RTN | **B=-0.007, SE=0.002, T=-4.235, p=0.000** | B=-0.005, SE=0.002, T=-2.093, p=0.036 |
| L_Hip | DAN | B=0.002, SE=0.001, T=2.048, p=0.041 | B=0.005, SE=0.001, T=3.445, p=0.001 |
| L_Hip | VAN | **B=-0.018, SE=0.001, T=-12.888, p=0.000** | **B=-0.010, SE=0.002, T=-4.792, p=0.000** |
| L_Hip | CPN | B=0.001, SE=0.001, T=1.629, p=0.103 | B=0.001, SE=0.001, T=0.749, p=0.454 |
| L_Hip | CON | **B=-0.007, SE=0.001, T=-5.493, p=0.000** | B=-0.002, SE=0.002, T=-1.160, p=0.246 |
| L_Hip | FPN | B=-0.005, SE=0.001, T=-3.266, p=0.001 | B=-0.005, SE=0.002, T=-2.442, p=0.015 |
| L_Hip | SN | B=-0.001, SE=0.001, T=-1.331, p=0.183 | B=-0.002, SE=0.002, T=-0.959, p=0.337 |
| L_Hip | DMN | **B=-0.007, SE=0.001, T=-7.292, p=0.000** | B=-0.003, SE=0.001, T=-2.413, p=0.016 |
| R_Hip | AN | **B=0.010, SE=0.002, T=6.091, p=0.000** | B=0.008, SE=0.002, T=3.225, p=0.001 |
| R_Hip | VN | **B=-0.011, SE=0.001, T=-7.429, p=0.000** | B=-0.005, SE=0.002, T=-2.317, p=0.021 |
| R_Hip | SMN-H | **B=-0.008, SE=0.001, T=-8.020, p=0.000** | B=-0.005, SE=0.001, T=-3.297, p=0.001 |
| R_Hip | SMN-M | B=-0.001, SE=0.001, T=-1.581, p=0.114 | B=-0.003, SE=0.001, T=-2.062, p=0.039 |
| R_Hip | RTN | **B=0.006, SE=0.001, T=6.195, p=0.000** | B=0.002, SE=0.001, T=1.323, p=0.186 |
| R_Hip | DAN | B=-0.001, SE=0.001, T=-0.849, p=0.396 | B=-0.004, SE=0.002, T=-2.536, p=0.011 |
| R_Hip | VAN | B=-0.003, SE=0.001, T=-3.907, p=0.000 | B=-0.001, SE=0.001, T=-0.964, p=0.335 |
| R_Hip | CPN | **B=-0.008, SE=0.001, T=-7.884, p=0.000** | B=-0.005, SE=0.002, T=-3.134, p=0.002 |
| R_Hip | CON | **B=-0.012, SE=0.001, T=-8.893, p=0.000** | **B=-0.008, SE=0.002, T=-4.111, p=0.000** |
| R_Hip | FPN | **B=-0.002, SE=0.001, T=-4.325, p=0.000** | B=-0.002, SE=0.001, T=-2.888, p=0.004 |
| R_Hip | SN | B=-0.003, SE=0.001, T=-2.068, p=0.039 | B=-0.004, SE=0.002, T=-1.951, p=0.051 |
| R_Hip | DMN | **B=-0.006, SE=0.001, T=-4.606, p=0.000** | B=-0.002, SE=0.002, T=-1.251, p=0.211 |

Statistics (including p-values) refer to models using the full sample, while bold highlights those connections that survived FDR corrections across split-half discovery and replication datasets (i.e., those connections highlighted as significant in Figure 1).

**Age and sex differences**

Supplemental analyses explored whether there were age or sex moderated associations between pubertal timing and corticolimbic connectivity. Linear mixed models were conducted for all set of analyses. Covariates included the fixed effects of sex, mean framewise displacement, outlier count, and participation following the start of the COVID19 pandemic (yes/no). Random effects of subject, family and site IDs were incorporated to account for the nested nature of the data. Results are presented in Table S3.

Table S3. *Associations between pubertal timing and corticolimbic connectivity, as a function of sex and age*

| **ROI** | **Network** | **Pubertal timing * Sex** | **Pubertal timing * Age** |
| --- | --- | --- | --- |
| L_Amy | AN | B=0.004, SE=0.002, T=1.699, p=0.089 | B=0.000, SE=0.000, T=1.281, p=0.200 |
| L_Amy | VN | B=-0.002, SE=0.002, T=-1.156, p=0.248 | B=0.000, SE=0.000, T=-1.233, p=0.218 |
| L_Amy | SMN-H | B=0.001, SE=0.002, T=0.624, p=0.533 | B=0.000, SE=0.000, T=1.322, p=0.186 |
| L_Amy | SMN-M | B=-0.006, SE=0.002, T=-2.389, p=0.017 | B=0.000, SE=0.000, T=-1.727, p=0.084 |
| L_Amy | RTN | B=0.004, SE=0.002, T=1.653, p=0.098 | B=0.000, SE=0.000, T=-0.505, p=0.613 |
| L_Amy | DAN | B=0.002, SE=0.001, T=1.360, p=0.174 | B=0.000, SE=0.000, T=1.729, p=0.084 |
| L_Amy | VAN | B=-0.001, SE=0.002, T=-0.405, p=0.685 | B=0.000, SE=0.000, T=0.747, p=0.455 |
| L_Amy | CPN | B=-0.002, SE=0.001, T=-2.680, p=0.007 | B=0.000, SE=0.000, T=-1.050, p=0.294 |
| L_Amy | CON | B=-0.009, SE=0.002, T=-4.373, p=0.000 | B=0.000, SE=0.000, T=-1.407, p=0.159 |
| L_Amy | FPN | B=-0.002, SE=0.001, T=-1.051, p=0.293 | B=0.000, SE=0.000, T=1.601, p=0.109 |
| L_Amy | SN | B=-0.001, SE=0.003, T=-0.503, p=0.615 | B=0.000, SE=0.000, T=0.121, p=0.904 |
| L_Amy | DMN | B=0.001, SE=0.002, T=0.376, p=0.707 | B=0.000, SE=0.000, T=-0.744, p=0.457 |
| R_Amy | AN | B=0.000, SE=0.002, T=-0.280, p=0.779 | B=0.000, SE=0.000, T=-0.194, p=0.846 |
| R_Amy | VN | B=-0.003, SE=0.002, T=-1.420, p=0.156 | B=0.000, SE=0.000, T=0.724, p=0.469 |
| R_Amy | SMN-H | B=-0.002, SE=0.002, T=-0.969, p=0.333 | B=0.000, SE=0.000, T=0.341, p=0.733 |
| R_Amy | SMN-M | B=0.002, SE=0.001, T=1.489, p=0.137 | B=0.000, SE=0.000, T=-0.762, p=0.446 |
| R_Amy | RTN | B=-0.001, SE=0.001, T=-1.310, p=0.190 | B=0.000, SE=0.000, T=0.208, p=0.835 |
| R_Amy | DAN | B=0.000, SE=0.002, T=0.116, p=0.908 | B=0.000, SE=0.000, T=0.724, p=0.469 |
| R_Amy | VAN | B=0.003, SE=0.001, T=2.268, p=0.023 | B=0.000, SE=0.000, T=-0.767, p=0.443 |
| R_Amy | CPN | B=-0.001, SE=0.002, T=-0.346, p=0.729 | B=0.000, SE=0.000, T=2.434, p=0.015 |
| R_Amy | CON | B=-0.006, SE=0.002, T=-2.671, p=0.008 | B=0.000, SE=0.000, T=-1.047, p=0.295 |
| R_Amy | FPN | B=-0.004, SE=0.002, T=-1.815, p=0.069 | B=0.000, SE=0.000, T=0.699, p=0.485 |
| R_Amy | SN | B=-0.004, SE=0.001, T=-2.635, p=0.008 | B=0.000, SE=0.000, T=-0.464, p=0.643 |
| R_Amy | DMN | B=0.000, SE=0.002, T=-0.114, p=0.910 | B=0.000, SE=0.000, T=-0.117, p=0.907 |
| L_Hip | AN | B=0.001, SE=0.001, T=1.335, p=0.182 | B=0.000, SE=0.000, T=1.939, p=0.053 |
| L_Hip | VN | B=-0.003, SE=0.002, T=-1.315, p=0.188 | B=0.000, SE=0.000, T=-1.364, p=0.173 |
| L_Hip | SMN-H | B=0.001, SE=0.002, T=0.439, p=0.661 | B=0.000, SE=0.000, T=2.004, p=0.045 |
| L_Hip | SMN-M | B=0.000, SE=0.002, T=-0.249, p=0.803 | B=0.000, SE=0.000, T=-0.968, p=0.333 |
| L_Hip | RTN | B=-0.001, SE=0.002, T=-0.483, p=0.629 | B=0.000, SE=0.000, T=-1.378, p=0.168 |
| L_Hip | DAN | B=0.004, SE=0.002, T=2.525, p=0.012 | B=0.000, SE=0.000, T=-0.559, p=0.576 |
| L_Hip | VAN | B=-0.007, SE=0.002, T=-3.313, p=0.001 | B=0.000, SE=0.000, T=-1.684, p=0.092 |
| L_Hip | CPN | B=0.000, SE=0.001, T=-0.045, p=0.964 | B=0.000, SE=0.000, T=-0.454, p=0.650 |
| L_Hip | CON | B=0.001, SE=0.002, T=0.328, p=0.743 | B=0.000, SE=0.000, T=0.563, p=0.573 |
| L_Hip | FPN | B=-0.003, SE=0.002, T=-1.137, p=0.256 | B=0.000, SE=0.000, T=-0.719, p=0.472 |
| L_Hip | SN | B=-0.002, SE=0.002, T=-1.211, p=0.226 | B=0.000, SE=0.000, T=-0.994, p=0.320 |
| L_Hip | DMN | B=-0.001, SE=0.001, T=-0.693, p=0.488 | B=0.000, SE=0.000, T=-0.758, p=0.448 |
| R_Hip | AN | B=0.005, SE=0.003, T=2.125, p=0.034 | B=0.000, SE=0.000, T=1.423, p=0.155 |
| R_Hip | VN | B=-0.001, SE=0.002, T=-0.417, p=0.676 | B=0.000, SE=0.000, T=-0.698, p=0.485 |
| R_Hip | SMN-H | B=-0.003, SE=0.002, T=-1.796, p=0.073 | B=0.000, SE=0.000, T=-1.072, p=0.284 |
| R_Hip | SMN-M | B=-0.003, SE=0.001, T=-1.942, p=0.052 | B=0.000, SE=0.000, T=-1.164, p=0.244 |
| R_Hip | RTN | B=-0.001, SE=0.001, T=-0.533, p=0.594 | B=0.000, SE=0.000, T=0.571, p=0.568 |
| R_Hip | DAN | B=-0.005, SE=0.002, T=-2.737, p=0.006 | B=0.000, SE=0.000, T=1.263, p=0.206 |
| R_Hip | VAN | B=-0.002, SE=0.001, T=-1.123, p=0.261 | B=0.000, SE=0.000, T=0.088, p=0.930 |
| R_Hip | CPN | B=-0.003, SE=0.002, T=-2.077, p=0.038 | B=0.000, SE=0.000, T=0.540, p=0.589 |
| R_Hip | CON | B=-0.003, SE=0.002, T=-1.588, p=0.112 | B=0.000, SE=0.000, T=-0.700, p=0.484 |
| R_Hip | FPN | B=-0.002, SE=0.001, T=-1.937, p=0.053 | B=0.000, SE=0.000, T=-0.840, p=0.401 |
| R_Hip | SN | B=-0.003, SE=0.002, T=-1.195, p=0.232 | B=0.000, SE=0.000, T=0.450, p=0.653 |
| R_Hip | DMN | B=-0.001, SE=0.002, T=-0.467, p=0.640 | B=0.000, SE=0.000, T=-0.660, p=0.509 |

Statistics (including p-values) refer to models using the full sample. There were no associations that survived FDR corrections across split-half discovery and replication datasets.

Table S4. *Mediation models*

|  | **Withdrawn Depression** | | | | **Rule-breaking Delinquency** | | | |
| --- | --- | --- | --- | --- | --- | --- | --- | --- |
|  | **Est** | **SE** | **CI*low*** | **CI*high*** | **Est** | **SE** | **CI*low*** | **CI*high*** |
| **L_Amy - CON** |  |  |  |  |  |  |  |  |
| CBCL ~ |  |  |  |  |  |  |  |  |
| Intercept | -0.544 | 0.097 | -0.735 | -0.354 | -0.585 | 0.112 | -0.807 | -0.365 |
| rsfMRI | -0.048 | 0.010 | -0.068 | -0.029 | 0.022 | 0.012 | -0.002 | 0.045 |
| pds_residual | 0.033 | 0.006 | 0.020 | 0.045 | 0.034 | 0.007 | 0.019 | 0.048 |
| sex | 0.431 | 0.049 | 0.337 | 0.530 | 0.216 | 0.045 | 0.127 | 0.303 |
| motion | 0.059 | 0.014 | 0.032 | 0.086 | 0.084 | 0.014 | 0.056 | 0.112 |
| outliers | -0.046 | 0.014 | -0.075 | -0.018 | 0.017 | 0.016 | -0.015 | 0.048 |
| covid | 0.002 | 0.060 | -0.116 | 0.118 | -0.040 | 0.073 | -0.183 | 0.102 |
| rsfMRI ~ |  |  |  |  |  |  |  |  |
| Intercept | 0.236 | 0.090 | 0.055 | 0.413 | 0.241 | 0.091 | 0.065 | 0.420 |
| pds_residual | -0.050 | 0.007 | -0.063 | -0.036 | -0.048 | 0.007 | -0.062 | -0.034 |
| sex | -0.009 | 0.017 | -0.041 | 0.026 | -0.008 | 0.017 | -0.043 | 0.025 |
| motion | -0.058 | 0.012 | -0.081 | -0.034 | -0.058 | 0.012 | -0.081 | -0.034 |
| outliers | 0.211 | 0.012 | 0.187 | 0.235 | 0.213 | 0.012 | 0.189 | 0.237 |
| covid | -0.224 | 0.068 | -0.358 | -0.090 | -0.230 | 0.069 | -0.368 | -0.093 |
| **R_Amy - CON** |  |  |  |  |  |  |  |  |
| CBCL ~ |  |  |  |  |  |  |  |  |
| Intercept | -0.560 | 0.097 | -0.750 | -0.373 | -0.569 | 0.108 | -0.781 | -0.353 |
| rsfMRI | -0.057 | 0.009 | -0.075 | -0.039 | -0.007 | 0.011 | -0.028 | 0.014 |
| pds_residual | 0.032 | 0.007 | 0.019 | 0.045 | 0.034 | 0.008 | 0.019 | 0.049 |
| sex | 0.431 | 0.048 | 0.337 | 0.528 | 0.213 | 0.044 | 0.126 | 0.299 |
| motion | 0.063 | 0.014 | 0.036 | 0.090 | 0.082 | 0.015 | 0.053 | 0.111 |
| outliers | -0.052 | 0.014 | -0.079 | -0.024 | 0.023 | 0.016 | -0.009 | 0.054 |
| covid | 0.019 | 0.059 | -0.098 | 0.136 | -0.055 | 0.072 | -0.198 | 0.084 |
| rsfMRI ~ |  |  |  |  |  |  |  |  |
| Intercept | 0.086 | 0.085 | -0.083 | 0.254 | 0.090 | 0.084 | -0.079 | 0.251 |
| pds_residual | -0.036 | 0.008 | -0.050 | -0.021 | -0.034 | 0.008 | -0.048 | -0.019 |
| sex | -0.004 | 0.017 | -0.038 | 0.029 | -0.004 | 0.017 | -0.039 | 0.030 |
| motion | -0.007 | 0.012 | -0.031 | 0.017 | -0.007 | 0.012 | -0.032 | 0.016 |
| outliers | 0.069 | 0.013 | 0.045 | 0.094 | 0.071 | 0.013 | 0.047 | 0.096 |
| covid | -0.092 | 0.071 | -0.231 | 0.050 | -0.098 | 0.071 | -0.233 | 0.044 |
| **R_Hip - CON** |  |  |  |  |  |  |  |  |
| CBCL ~ |  |  |  |  |  |  |  |  |
| Intercept | -0.562 | 0.100 | -0.763 | -0.362 | -0.585 | 0.111 | -0.803 | -0.369 |
| rsfMRI | -0.032 | 0.010 | -0.051 | -0.012 | 0.039 | 0.012 | 0.017 | 0.062 |
| pds_residual | 0.034 | 0.006 | 0.021 | 0.047 | 0.034 | 0.007 | 0.019 | 0.048 |
| sex | 0.431 | 0.049 | 0.336 | 0.526 | 0.214 | 0.044 | 0.129 | 0.298 |
| motion | 0.059 | 0.014 | 0.032 | 0.086 | 0.083 | 0.014 | 0.055 | 0.111 |
| outliers | -0.049 | 0.014 | -0.078 | -0.022 | 0.016 | 0.016 | -0.015 | 0.047 |
| covid | 0.020 | 0.060 | -0.098 | 0.137 | -0.043 | 0.074 | -0.187 | 0.102 |
| rsfMRI ~ |  |  |  |  |  |  |  |  |
| Intercept | 0.217 | 0.097 | 0.028 | 0.408 | 0.219 | 0.097 | 0.029 | 0.408 |
| pds_residual | -0.030 | 0.007 | -0.044 | -0.016 | -0.028 | 0.007 | -0.042 | -0.014 |
| sex | -0.006 | 0.017 | -0.040 | 0.027 | -0.006 | 0.017 | -0.040 | 0.028 |
| motion | -0.039 | 0.012 | -0.062 | -0.015 | -0.039 | 0.012 | -0.062 | -0.015 |
| outliers | 0.142 | 0.012 | 0.118 | 0.165 | 0.144 | 0.012 | 0.120 | 0.167 |
| covid | -0.209 | 0.068 | -0.344 | -0.077 | -0.211 | 0.067 | -0.340 | -0.078 |
| **L_Hip - VAN** |  |  |  |  |  |  |  |  |
| CBCL ~ |  |  |  |  |  |  |  |  |
| Intercept | -0.563 | 0.099 | -0.755 | -0.366 | -0.575 | 0.109 | -0.790 | -0.368 |
| rsfMRI | -0.033 | 0.009 | -0.051 | -0.015 | 0.017 | 0.011 | -0.005 | 0.038 |
| pds_residual | 0.033 | 0.006 | 0.020 | 0.045 | 0.034 | 0.008 | 0.019 | 0.049 |
| sex | 0.432 | 0.049 | 0.335 | 0.528 | 0.215 | 0.044 | 0.130 | 0.304 |
| motion | 0.061 | 0.014 | 0.034 | 0.089 | 0.083 | 0.015 | 0.054 | 0.112 |
| outliers | -0.057 | 0.014 | -0.086 | -0.030 | 0.021 | 0.016 | -0.011 | 0.054 |
| covid | 0.023 | 0.059 | -0.092 | 0.137 | -0.049 | 0.072 | -0.189 | 0.093 |
| rsfMRI ~ |  |  |  |  |  |  |  |  |
| Intercept | 0.372 | 0.085 | 0.202 | 0.536 | 0.372 | 0.084 | 0.207 | 0.539 |
| pds_residual | -0.036 | 0.008 | -0.051 | -0.021 | -0.035 | 0.008 | -0.050 | -0.020 |
| sex | -0.009 | 0.017 | -0.042 | 0.025 | -0.008 | 0.017 | -0.043 | 0.025 |
| motion | 0.010 | 0.013 | -0.015 | 0.035 | 0.012 | 0.012 | -0.012 | 0.036 |
| outliers | -0.020 | 0.013 | -0.045 | 0.005 | -0.021 | 0.012 | -0.045 | 0.003 |
| covid | -0.381 | 0.072 | -0.521 | -0.241 | -0.383 | 0.071 | -0.518 | -0.243 |

pds_residual: pubertal timing; race/ethnicity: compared to Black/African American, covid: pre vs post start of pandemic, motion: mean framewise displacement

**Controlling for race/ethnicity and INR**

Supplemental analyses examined whether incorporation of race/ethnicity and INR as additional covariates influenced the main findings. The addition of these covariates into linear mixed models did not change the significance of associations between pubertal timing and connectivity of left amygdala-CON (p < 0.001), right amygdala-CON (p = 0.008) or left hippocampus-VAN (p < 0.001). However, the association between timing and right hippocampus-CON was no longer significant (p = 0.112). Inclusion of these covariates into mediation models did not change the significance of indirect pathways between pubertal timing and withdrawn depression via amygdala-CON or left hippocampus-VAN. However, indirect pathways involving the right hippocampus-CON (to both depression and delinquency) were no longer significant (see Figure S2).

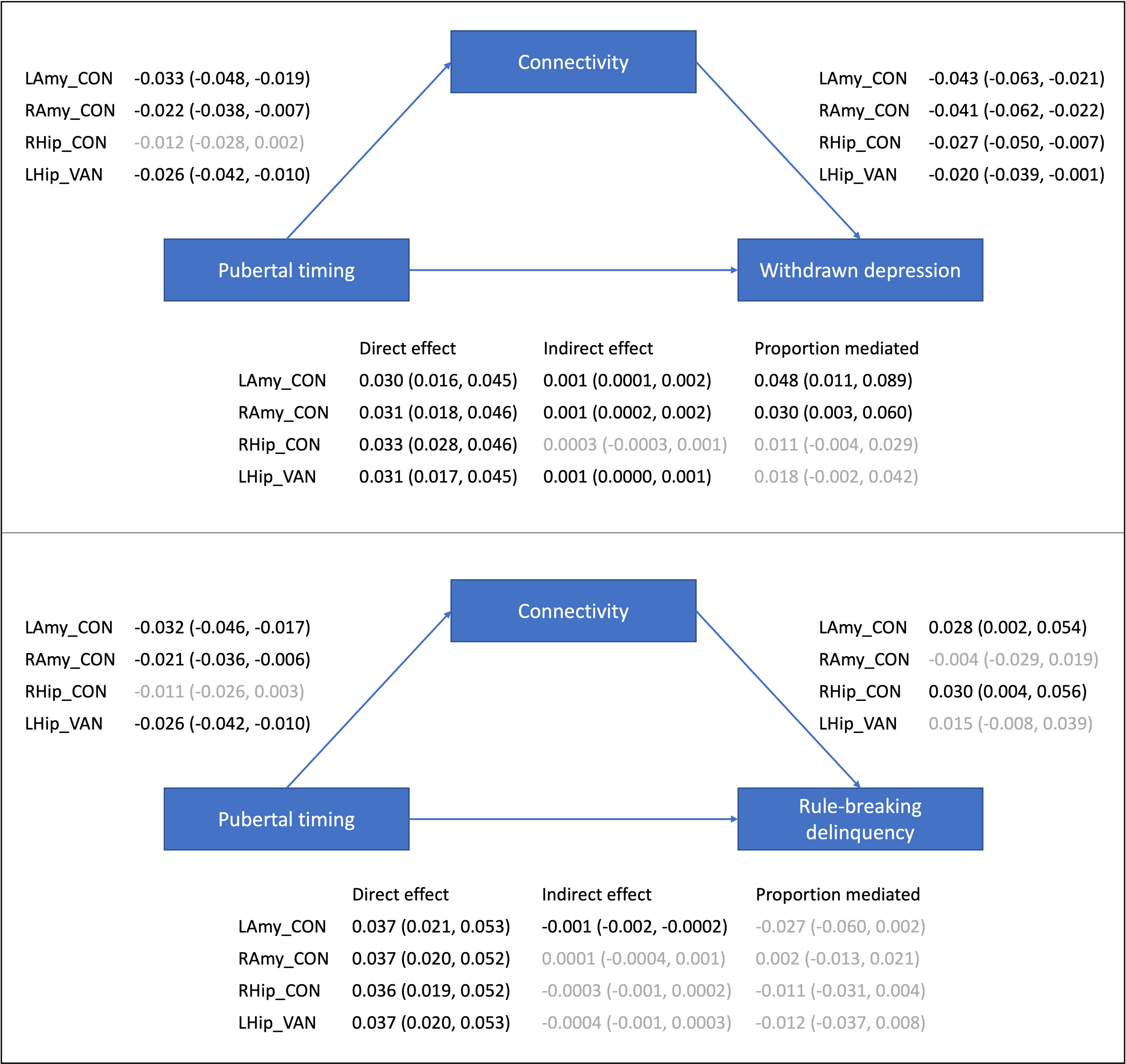

*Figure S2.* Mediation models examining indirect pathways between pubertal timing and mental health problems via resting-state corticolimbic connectivity, while controlling for race/ethnicity and INR (estimates and 95% CIs presented).
